# Spatio-temporal surveys of the brown meagre *Sciaena umbra* using passive acoustics for management and conservation

**DOI:** 10.1101/2020.06.03.131326

**Authors:** Lucia Di Iorio, Patrick Bonhomme, Noëmie Michez, Bruno Ferrari, Alexandra Gigou, Pieraugusto Panzalis, Elena Desiderà, Augusto Navone, Pierre Boissery, Julie Lossent, Benjamin Cadville, Marie Bravo-Monin, Eric Charbonnel, Cédric Gervaise

## Abstract

Conservation of exploited fish populations is a priority for environmental managers. Spatio-temporal knowledge on reproductive sites is mandatory for species and habitat conservation but is often difficult to assess, particularly over vast geographic areas. Regular and long-term standardized surveys are necessary to identify reproductive sites, assess population trends and their distribution. Here we emphasize the utility of Passive Acoustic Monitoring (PAM) for the survey and management of a depleted vulnerable Mediterranean fish species, the brown meagre, *Sciaena umbra*. Acoustic surveys of reproductive calls were conducted combining 1) spatial data from standardized surveys within three MPAs and from 49 unprotected sites throughout the Northwestern Mediterranean basin, as well as 2) temporal data from a two-year-long survey at a presumed spawning location. The MPA surveys, which rapidly scanned ~30-50 km of the rocky coastlines per MPA, unveiled maps of distribution and reproductive activity of the brown meagre, including potential spawning sites. They were also effective in emphasizing effects linked to management actions: Full-protection zones had a higher number of vocalizations (70% of the listening sites) compared to less protected zones (30% of the sites) or sites outside MPAs (45% of the sites). This was also reflected in the number of singers that was generally low (< 3 individuals) in less protected zones and outside MPAs, implying lower fish densities. Highest calling aggregations were observed in potential spawning areas that represented only 0.04% of all listening sites, and were almost all in older, fully protected MPAs, which thus play a key role for fish stock recovery. The two-year survey revealed a 5-month reproductive season (from May to October) with a strong positive correlation between calling activity and temperature. Overall this study confirms the role of PAM as an efficient, replicable and standardized non-invasive method for population management that can identify functional sites and key protection zones, provide valuable information on reproduction, spatial and temporal occurrence, but also on population trends and climate-driven changes.

**Highlights:**

- Monitoring of threatened species and their key habitats is critical for environmental managers.
- Management requires methods to assess population trends at large spatial and temporal scales.
- Passive acoustics (PA) is efficient in mapping and monitoring vulnerable fish species.
- Distribution, reproductive sites and population dynamics can be assessed over vast geographical areas.
- We show the utility of PA to identify key conservation zones and assess effects of management actions.

## 1 INTRODUCTION

Knowledge of spawning habitats and breeding sites is essential for the conservation of exploited fish stocks. Recording when and where a fish species is reproducing provides pivotal data on key habitats and allows to better manage both vulnerable species and their critical habitats (Luczkovich et al., 2008b). Identification, mapping and monitoring of functional fish habitats is therefore a priority for environmental managers of Marine Protected Areas (MPA), in particular with regard to endangered and over-exploited species (Harmelin and Ruitton, 2006; Luczkovich et al., 2008a). Regular and long-term surveys of target species at large spatial scales are necessary to evaluate population trends, distribution, recovery, and the efficiency of protection measures (Guidetti et al., 2014a; Harmelin-Vivien et al., 2015). However, although very effective, traditional methods based on visual surveys are logistically difficult to implement at high-frequency intervals, over wide areas and for nocturnally active organisms (Harmelin-Vivien et al. 2015). Management programs would therefore benefit from complementary standardized and replicable methods allowing to cover long distances over short time periods and to monitor key sites and key species continuously, over the long-term with limited human investment.

Passive acoustic monitoring (PAM) has proven to be a powerful tool for both conservation and fishery science (e.g., Gannon, 2008; Luczkovich et al., 2008a; Mann et al., 2008). It is a non-invasive flexible listening technique that involves the use of hydrophones to record the collection of sounds emanated from an environment (i.e., the soundscape). PAM exploits the sounds emitted by fish mainly for communication (e.g., Amorim, 2006; Ladich, 2015) to help identifying species occurrence, spawning areas, cryptic species, follow reproductive activity and population turnover, and to appraise diversity (Desiderà et al., 2019; Lobel, 2002; Luczkovich et al., 2008b; Picciulin et al., 2018). Despite this, and the fact that a variety of commercial and vulnerable fish species produce sounds, there is a limited number of studies adopting PAM for large-scale mapping of protected fish species and their critical habitats in the Mediterranean Sea, and they all concern the brown meagre *Sciaena umbra* Linnaeus 1758 (Bonacito et al., 2002; Parmentier et al., 2018; Picciulin et al., 2012).

The brown meagre is an iconic demersal fish species of Mediterranean coastal habitats listed in the Annexe III (Protected Fauna Species) of the Barcelona and Berne Conventions. Globally, this species is classified as Near Threatened (Chao, 2015) and as vulnerable in the Mediterranean Sea by the IUCN (Abdul Malak et al., 2011; Bizsel et al., 2011). *S. umbra* is highly vulnerable to fishing due to its aggregative behaviour and easily accessible shallow-water habitat (Lloret et al., 2008). With a population decline in the Mediterranean Sea of approximately 70% between 1980 and 2005 (Bizsel et al., 2011), the brown meagre falls within the categories of seriously depleted and collapsed fish species. This is why in 2013, France adopted a moratorium banning spear and recreational fishing of the brown meagre, which was renewed in December 2018. *S. umbra* is also of high ecological and conservation value as it is considered a useful bioindicator (Mouillot et al., 2002). Its presence is indicative of high environmental quality and community richness (García-Rubies et al., 2013; Mouillot et al., 2002; Picciulin et al., 2013), and it is very sensitive to the effect of protection measures (Guidetti et al., 2014b; Harmelin-Vivien et al., 2015b; Harmelin et al., 1995). Although there is conspicuous increase in its abundance in no-take integral reserves (Francour, 1991; García-Rubies et al., 2013; Harmelin, 2013), outside most protected areas, the brown meagre is frequently targeted (Lloret et al., 2008; Morales-Nin et al., 2005) and found to be rare or absent (Harmelin-Vivien et al., 2015b; Letourneur et al., 2003). Appropriate monitoring of *S. umbra* populations and their trends should involve the combination of extensive large-scale observations with regular long-term monitoring of key sites within and outside MPAs (Harmelin & Ruitton, 2007; Harmelin, 2013). This would allow appropriate assessment of species distribution as well as its spatial extension and to identify and follow functional sites (e.g., feeding, reproduction, spawning).

*S. umbra* shows strong site-fidelity particularly during the autumn and winter months (Alós and Cabanellas-Reboredo, 2012; Harmelin and Ruitton, 2006), but there is a lack of knowledge on the reproductive movements occurring during the summer. As many other sciaenid species, the brown meagre forms breeding aggregations (Bono et al., 2001; Grau et al., 2009; Ragonese et al., 2002). Locations of aggregations and spawning sites need to be better documented in order to be protected and managed (Harmelin-Vivien et al., 2015). During the spawning period, i.e., May until August (Chauvet, 1991, Grau et al., 2009), *S. umbra* males produce sounds (Parmentier et al., 2017, Picciulin et al., 2012). In other Sciaenids, these types of vocalizations have been shown to serve to aggregate individuals, attract mates to leks (Gilmore, 2002; Parsons et al., 2009), advertise male readiness to spawn, and synchronise gamete release (Connaughton & Taylor 1995, Lobel, 1992). *S. umbra* vocalizations are loud percussive sounds produced by a sonic muscles associated to the males’ swim bladder (Dijkgraaf, 1947, Parmentier et al., 2017). These percussive sounds can either be produced as irregular sounds (I-calls), or in long regular stereotyped sequences of calls (R-calls) (Picciulin et al. 2012) emitted for several hours at nighttime (Fig. 1). In Sciaenids these R-calls represent reproductive displays. Their occurrence in high densities forms so-called choruses, which are indicative of spawning aggregations (Guest and J., 1978; Picciulin et al. 2012, Parsons et al., 2009, Luczkovich et al., 2008b; Mok and Gilmore, 1983). Therefore, sound production has the potential to serve 1) as a proxy for *S. umbra’s* reproductive activity, 2) to identify nocturnal individual occurrence as well as breeding areas and 3) to locate and monitor aggregations and spawning sites over extended time periods (Ramcharitar et al., 2006). Despite the potential of PAM to fill some of the knowledge gaps about the brown meagre’s ecology and reproductive behaviour and a few pioneering studies in the Gulf of Venice (Italy) using PAM for spatial monitoring (Bonacito et al., 2002; Colla et al., 2018; Picciulin et al., 2013b), there are no studies reporting the application and validation of this technique by environmental managers. The aims of this study were to underline the efficiency of PAM for the survey and management of *S. umbra* and its potential spawning grounds within and outside MPAs by mapping it’s acoustic activity 1) along the coastline of three wide-ranging MPAs, 2) at large spatial scales (Northwestern Mediterranean), and 3) over two years in a no-take reserve.

**Figure 1.**
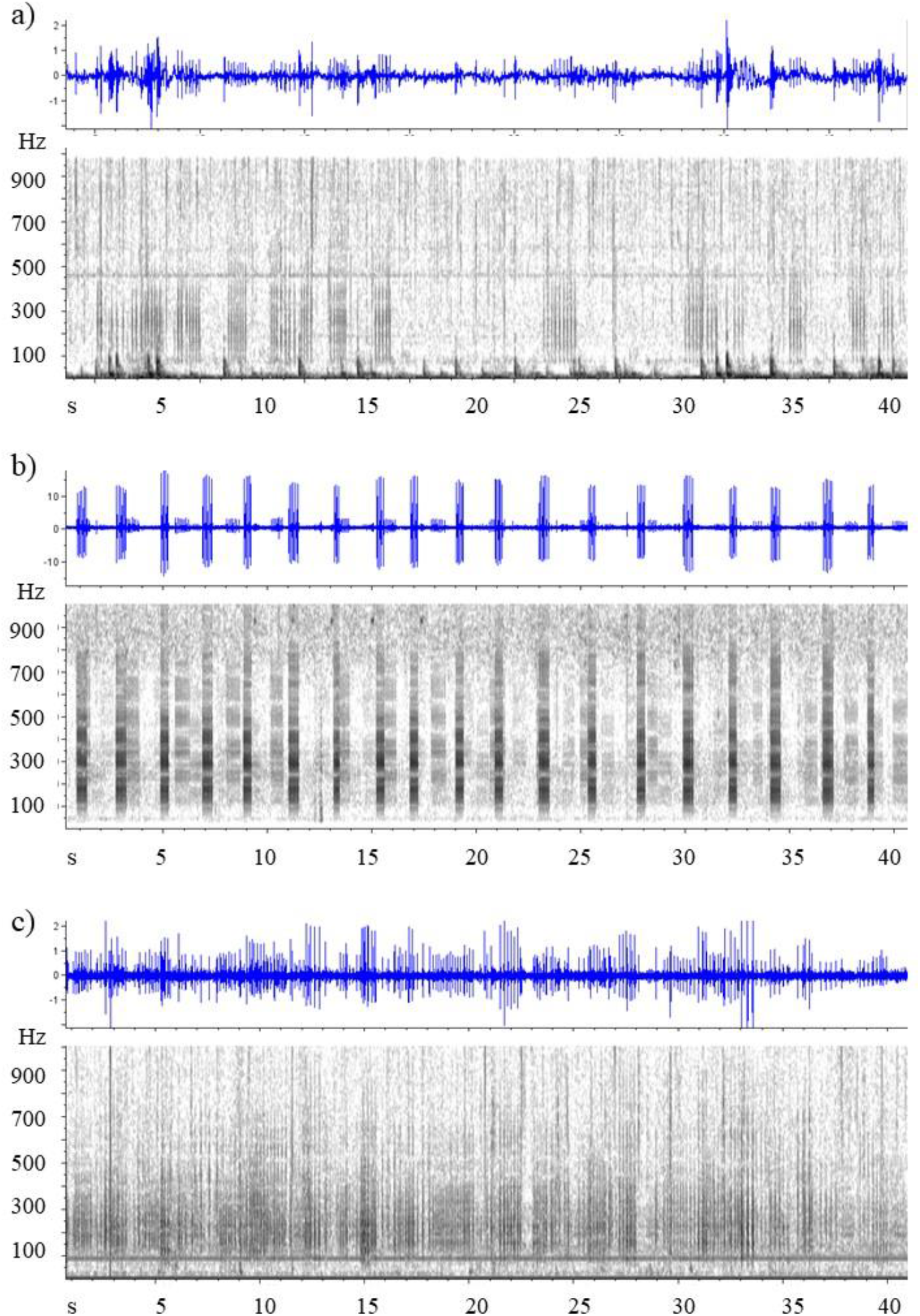
Oscillograms (top) and spectrograms (bottom) of *S. umbra* call categories. a) Irregular I-calls, b) series of stereotyped R-calls, and c) choruses, as described by Picciulin et al. 2012.

## 2 METHODS

### 2.1 Spatial sampling within three MPAs

#### 2.1.1 Study sites

The study sites were three vast MPAs in the Northwestern Mediterranean Sea: The MPA of the Calanques National Park (CNP), the Natural Marine Park of the Gulf of Lion (NMPGL), both in France, and the MPA of Tavolara-Punta Coda Cavallo (TPCC) in Sardinia, Italy (Fig. 2). The CNP is the youngest French national park established in 2012 and situated between Marseille and La Ciotat (Fig. 2). The AMP comprises a 435 km^2^ area that includes seven no-take zones (NTZ, no fishing) and one reinforced protection zone (RPZ) that allows yet regulates artisanal and recreational fishing activity. The NMPGL is also a young MPA, established in 2011 with a vast surface of 4000 km^2^ along 100 km coastline. The NMPGL includes the Natural Marine Reserve of Cerbère-Banyuls (MRCB), which is the oldest marine reserve in France created in 1974, measuring 6.50 km^2^ and consisting of a RPZ where only recreational fishing and diving are allowed and regulated and a fully protected no-take and no-access zone (FPZ) (Fig. 2). Finally, the TPCC MPA in Northeastern Sardinia, Italy, established in 1997 but effectively managed around 2003-04, comprises 76 km of coastline and a surface area of 153.57 km^2^. Three types of zones with different levels of protection have been established: (1) A zones, corresponding to FPZs (i.e., no-take and no-access); (2) B zones, corresponding to RPZ, where only licensed local artisanal fishing and diving are allowed; (3) C zones, corresponding to general protection zones (PZ), where both artisanal and recreational fishing are authorized as well as diving (Fig. 2).

**Figure 2.**
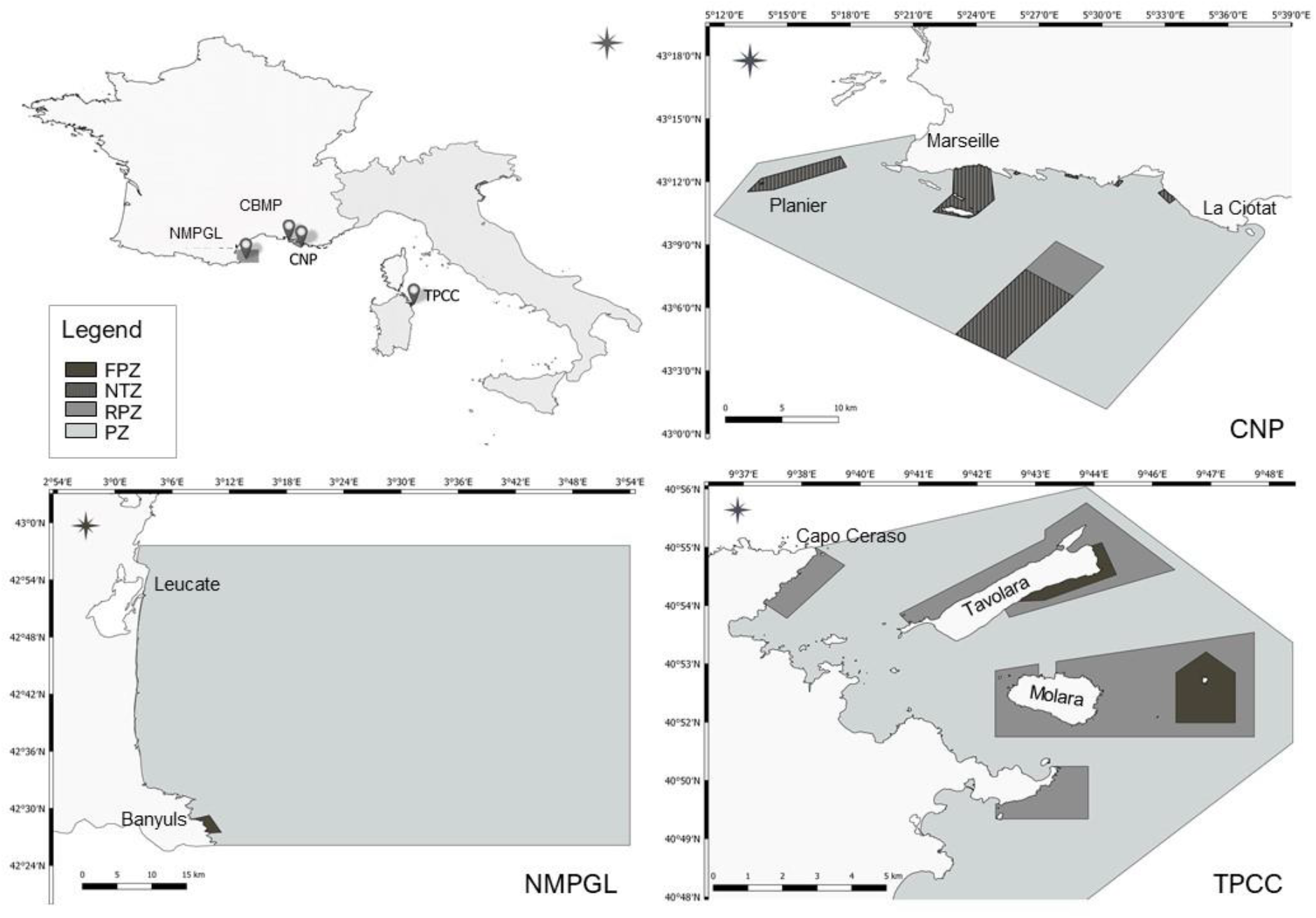
Maps of the MPAs where drifting surveys were conducted. Top left panel: France and Italy with the locations of the four MPAs. Other panels: Maps of the three MPAs in which *S. umbra* distribution was assessed using drifting devices. The different protection levels are indicated by different colours described in the legend: FPZ = Full Protection Zone (=Reserve = A zone), NTZ = No-Take-Zone (only in PNC), RPZ = Reinforced Protection Zone (= B zone), PZ = Protection Zone (= C zone). CNP: Calanques National Park, NMPGL: Natural Marine Park of the Gulf of Lion with the Marine Reserve of Cerbère-Banyuls (dark grey), TPCC: MPA of Tavolara-Punta Coda Cavallo.

#### 2.1.2 Acoustic data acquisition and recording protocol

Spatial mapping of *S. umbra* based on their vocalisations has been proposed and successfully adopted in the Gulf of Venice (Italy) (Bonacito et al., 2002; Colla et al., 2018; Picciulin et al., 2013). In this study sound recordings were acquired using two or three simultaneously deployed drifting devices equipped with an EA-SDA14 compact autonomous recorder (RTSys^®^, France) and a HTI-92-WB (High Tech Inc.^®^) hydrophone with a flat sensitivity equal to −155 ± 3 dB re 1 V μPa-1 between 5 and 50 kHz. Acoustic recordings were sampled continuously at a rate of 78 kHz and a 24-bit resolution. The hydrophone was located between 5 m and 9 m below the sea surface. Positions were defined based on the mean propagation range of *S. umbra* calls estimated using reported source levels (Codarin et al. 2009, Parmentier et al. 2017) and assuming cylindrical spreading loss. Consequently, maximal inter-recording position-distance was set at 600m. Priority and secondary zones for the survey were selected by the MPA’s agents based on habitat, protection level and existing knowledge. Deployments were performed using a standardized protocol: Along rocky coastlines, the devices were deployed within maximum 200m from the coast, one after the other at 200 to 600m distance, with water depths ranging between 8 and 20m. After deployment of the drifting recorders at a target positions, the boat moved 200 m away and stopped the engine in order to avoid noise interference. After 7 min of recording, each device was recovered. At each deployment and recovery, GPS time and position were noted. All drifts were conducted during the summer, from 7 p.m. to 12 p.m., which includes the period of maximal vocalizing activity of the brown meagre (Parmentier et al. 2017; Picciulin et al. 2013) (Table 1).

**Table 1.** Summary table of the different survey types and the recording sites. For precautionary reasons with regard to illegal fishing, the coordinates of the positions are not provided.

| ID | Site | Habitat | Survey | Survey dates | Year | *S. umbra* | Recorders | Recorder support | Recording stations |
| --- | --- | --- | --- | --- | --- | --- | --- | --- | --- |
|  | CNP | rock | MPA | 26-28/06/2018 | 2018 | 1 | 3 | floating device | 72 |
|  | NMPGL | rock | MPA | 24-27/06 & 29/08/2019 | 2019 | 1 | 3 | floating device | 96 |
|  | TPCC | rock | MPA | 22-24/07/2019 | 2019 | 1 | 2 | floating device | 54 |
| 1 | Banuyls | coralligenous | NW Med Sea | 2016/06/21 | 2016 | 0 | 1 | bottom mooring | 1 |
| 1 | Banuyls | seagrass | NW Med Sea | 2016/06/21 | 2016 | 1 | 1 | bottom mooring | 1 |
| 2 | Agde | coralligenous | NW Med Sea | 2016/06/20 | 2016 | 1 | 1 | bottom mooring | 1 |
| 3 | Grand Travers | coralligenous | NW Med Sea | 2016/06/17 | 2016 | 0 | 1 | bottom mooring | 1 |
| 4 | Ciotat | seagrass | NW Med Sea | 2016/06/02 | 2016 | 0 | 1 | bottom mooring | 1 |
| 4 | Ciotat | coralligenous | NW Med Sea | 2016/06/02 | 2016 | 1 | 1 | bottom mooring | 1 |
| 5 | Sicie | seagrass | NW Med Sea | 2016/06/03 | 2016 | 0 | 1 | bottom mooring | 1 |
| 5 | Sicie | coralligenous | NW Med Sea | 2016/06/03 | 2016 | 1 | 1 | bottom mooring | 1 |
| 6 | Giens | coralligenous | NW Med Sea | 2016/06/04 | 2016 | 1 | 1 | bottom mooring | 1 |
| 7 | Ribaud | seagrass | NW Med Sea | 2016/06/04 | 2016 | 1 | 1 | bottom mooring | 1 |
| 8 | Bormes | coralligenous | NW Med Sea | 2016/06/05 | 2016 | 0 | 1 | bottom mooring | 1 |
| 8 | Bormes | seagrass | NW Med Sea | 2016/06/05 | 2016 | 1 | 1 | bottom mooring | 1 |
| 9 | Cap Lardier | coralligenous | NW Med Sea | 2016/06/06 | 2016 | 0 | 1 | bottom mooring | 1 |
| 9 | Cap Lardier | seagrass | NW Med Sea | 2016/06/06 | 2016 | 1 | 1 | bottom mooring | 1 |
| 10 | Bonneau | seagrass | NW Med Sea | 2016/06/08 | 2016 | 1 | 1 | bottom mooring | 1 |
| 10 | Bonneau | coralligenous | NW Med Sea | 2016/06/08 | 2016 | 1 | 1 | bottom mooring | 1 |
| 11 | Cap Roux | seagrass | NW Med Sea | 2016/06/09 | 2016 | 0 | 1 | bottom mooring | 1 |
| 11 | Cap Roux | coralligenous | NW Med Sea | 2016/06/09 | 2016 | 0 | 1 | bottom mooring | 1 |
| 12 | Golfe Juan | seagrass | NW Med Sea | 2016/06/10 | 2016 | 0 | 1 | bottom mooring | 1 |
| 12 | Golfe Juan | coralligenous | NW Med Sea | 2016/06/10 | 2016 | 0 | 1 | bottom mooring | 1 |
| 13 | Bacon | coralligenous | NW Med Sea | 2016/06/11 | 2016 | 0 | 1 | bottom mooring | 1 |
| 13 | Bacon | seagrass | NW Med Sea | 2016/06/11 | 2016 | 0 | 1 | bottom mooring | 1 |
| 14 | Nice | coralligenous | NW Med Sea | 2016/06/12 | 2016 | 0 | 1 | bottom mooring | 1 |
| 14 | Nice | seagrass | NW Med Sea | 2016/06/13 | 2016 | 1 | 1 | bottom mooring | 1 |
| 15 | Eze | coralligenous | NW Med Sea | 2016/06/13 | 2016 | 0 | 1 | bottom mooring | 1 |
| 16 | Cap Martin | coralligenous | NW Med Sea | 2016/06/15 | 2016 | 0 | 1 | bottom mooring | 1 |
| 17 | Maccinagio | seagrass | NW Med Sea | 2017/06/02 | 2017 | 1 | 1 | bottom mooring | 1 |
| 17 | Maccinagio | coralligenous | NW Med Sea | 2017/06/02 | 2017 | 1 | 1 | bottom mooring | 1 |
| 18 | Bastia | coralligenous | NW Med Sea | 2017/05/31 | 2017 | 0 | 1 | bottom mooring | 1 |
| 18 | Bastia | seagrass | NW Med Sea | 2017/05/31 | 2017 | 0 | 1 | bottom mooring | 1 |
| 19 | Tarco | coralligenous | NW Med Sea | 2017/05/30 | 2017 | 0 | 1 | bottom mooring | 1 |
| 19 | Tarco | seagrass | NW Med Sea | 2017/05/30 | 2017 | 1 | 1 | bottom mooring | 1 |
| 20 | Rondinara | coralligenous | NW Med Sea | 2017/05/29 | 2017 | 0 | 1 | bottom mooring | 1 |
| 20 | Rondinara | seagrass | NW Med Sea | 2017/05/29 | 2017 | 1 | 1 | bottom mooring | 1 |
| 21 | Murtoli | seagrass | NW Med Sea | 2017/05/27 | 2017 | 1 | 1 | bottom mooring | 1 |
| 22 | Parata | seagrass | NW Med Sea | 2017/06/12 | 2017 | 1 | 1 | bottom mooring | 1 |
| 22 | Parata | coralligenous | NW Med Sea | 2017/06/12 | 2017 | 1 | 1 | bottom mooring | 1 |
| 23 | Cappo Rossu | seagrass | NW Med Sea | 2017/06/10 | 2017 | 1 | 1 | bottom mooring | 1 |
| 23 | Cappo Rossu | coralligenous | NW Med Sea | 2017/06/10 | 2017 | 0 | 1 | bottom mooring | 1 |
| 24 | Focolara | coralligenous | NW Med Sea | 2017/06/09 | 2017 | 0 | 1 | bottom mooring | 1 |
| 24 | Focolara | seagrass | NW Med Sea | 2017/06/09 | 2017 | 1 | 1 | bottom mooring | 1 |
| 25 | Calvi | seagrass | NW Med Sea | 2017/06/05 | 2017 | 1 | 1 | bottom mooring | 1 |
| 25 | Calvi | coralligenous | NW Med Sea | 2017/06/05 | 2017 | 1 | 1 | bottom mooring | 1 |
| 26 | Agriates | coralligenous | NW Med Sea | 2017/06/03 | 2017 | 0 | 1 | bottom mooring | 1 |
| 26 | Agriates | seagrass | NW Med Sea | 2017/06/03 | 2017 | 1 | 1 | bottom mooring | 1 |
| CBMP | Côte Bleue | seagrass | NW Med Sea | 2016/05/30 | 2016 | 0 | 1 | bottom mooring | 1 |
| 27 | Cala Gonone | rock | NW Med Sea | several months | 2017 | 1 | 1 | bottom mooring | 1 |
| CBMP | Côte Bleue | rock | NW Med Sea | yearround | 2017/2018 | 1 | 1 | bottom mooring | 1 |
| TPCC | Tavolara | coralligenous | NW Med Sea | several months | 2017 | 1 | 1 | bottom mooring | 1 |

### 2.2 Spatial sampling across the Northwestern Mediterranean basin

Sound recordings from 49 stations of the CALME acoustic network along the French Western Mediterranean coast established by the RMC Water Agency and the CHORUS Institute (Gervaise et al. 2018) were used; 22 from *Posidonia oceanica* meadows, and 25 from coralligenous reefs, and two from rocky reefs obtained in summer 2016 and 2017 (Table 1). The CALME network was not designed for *S. umbra* monitoring but is dedicated to key habitat and noise surveillance within the scope of EU Water Framework Directive (2000/60/EC) and the Marine Strategy Framework Directive (2008/56/EC). The same recording equipment (hydrophones and autonomous recorders) was deployed as for the drifting devices. The recorders were bottom moored with the hydrophone at 1 m from the seafloor (Fig. 3). Recorders acquired sounds continuously at a 78 kHz sampling rate and 24-bit resolution. For 46 of the 50 recording stations, at each recording date, the recorder was submerged in the afternoon and recovered the next morning for a duration of at least 14 hours. Recordings were made during night-time because the brown meagre, as most temperate fishes, vocalises at night (Parmentier et al., 2017; Parsons et al., 2016; Picciulin et al., 2012). In four sites, recordings were made over more than 24h.

**Figure 3.**
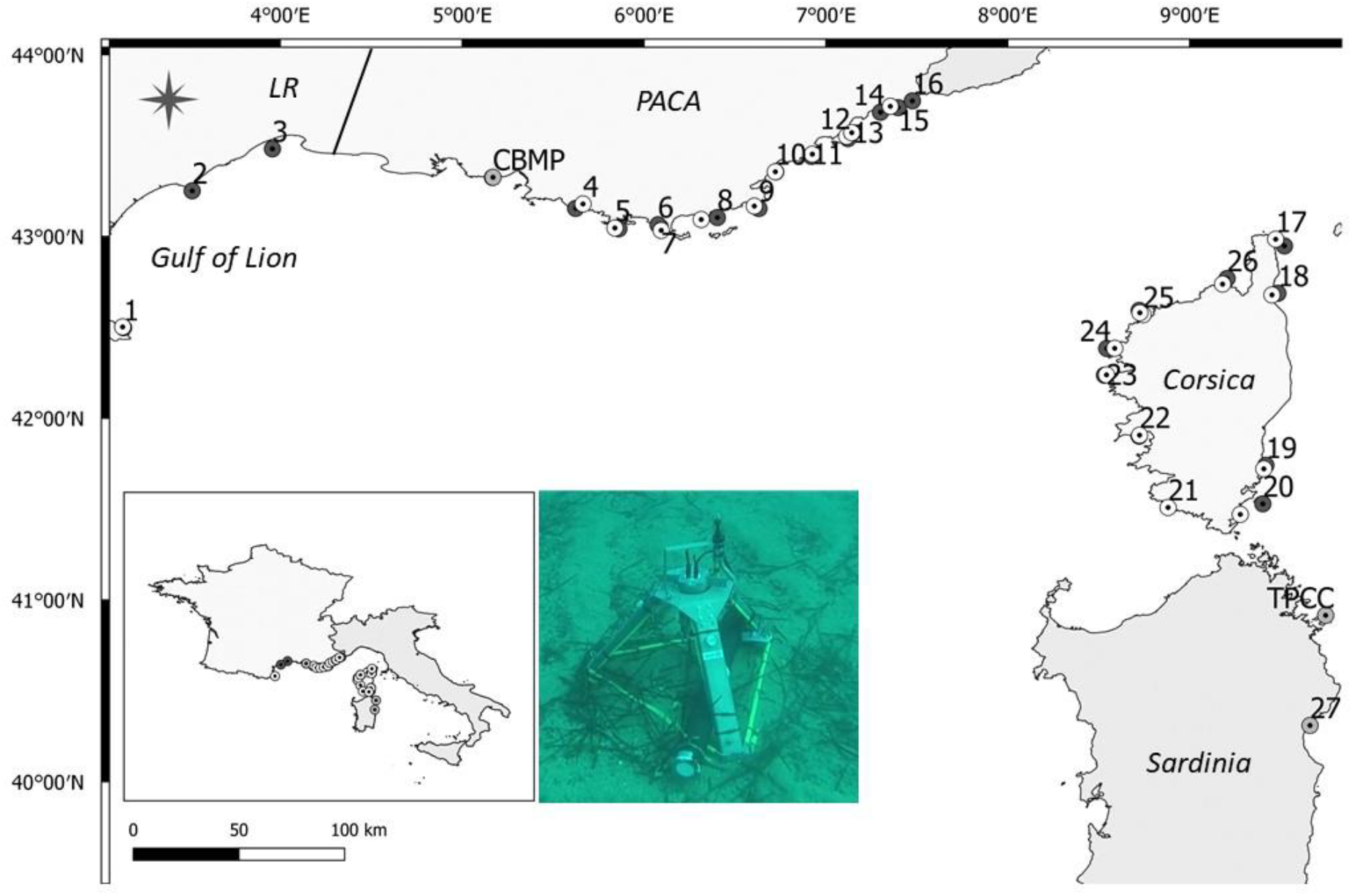
Map of the 49 recording sites of the large-scale monitoring program CALME and the long-term recording site in the Côte Bleue Marine Park (CBMP). Dark grey circles indicate coralligenous reef recording sites, bright grey circle indicate rocky long-term recording stations, white circles are seagrass meadow recording sites. The picture illustrates the mooring system with the recording device. The locations corresponding to the numbers are listed in table 1. *PACA*: Provence Alpes Côte d’Azur region, *LR*: Languedoc-Roussillon region.

### 2.3 Long-term data collection

One long-term recording from the CALME network situated in typical *S. umbra* habitat was used to assess annual trends in acoustic presence. The recording was obtained from the Réserve de Couronne in the Côte Bleue Marine Park (CBMP), near Marseille, France (43° 19’ 32”, 5° 10’ 11”). This reserve was created in 1996 and comprises 210 ha. It is a FPZ, where all kind of fishing activities, mooring and scuba diving are prohibited. An EA-SDA14 compact autonomous recorder (RTSys^®^, France) and an HTI-92-WB (High Tech Inc.^®^) with the same characteristics described for the drifting device was used but equipped with an external battery pack for long-term deployments. Data were recorded continuously and year-round in 2017 and 2018. The recorder was placed on a spawning site of seabreams (*Sparus aurata*) and seabass (*Dicentrarchus labrax*) known to regularly host brown meagres. The recording site was situated near an artificial reef and a natural coralligenous reef, next to a patchy *Posidonia oceanica* meadow, at a depth of 25 m. Bottom temperature profiles and trends were obtained from a temperature probe installed nearby by the park’s agents since 1998.

### 2.4 Data processing

All data was first down-sampled to 4kHz. On the one hand, data acquired during the acoustic small-scale spatial surveys within the MPAs and those obtained from the large-scale CALME network were visually inspected using Raven PRO 1.5 (The Cornell Lab of Ornithology; Hanning window, LFFT = 512, overlap: 50%). All recordings from the MPA surveys were inspected for presence/absence of I-calls, R-calls and choruses, as the presence of these different call types may indicate functional differences of sites. I-calls are characterized by variable numbers of pulses with no fixed call repetition rate. In contrast, R calls are composed of 5-6 pulses repeated in long regular stereotyped sequences (Picciulin et al. 2012) (Fig. 1). They are mainly produced between 5 p.m. and 12 p.m. with a peak between 8 p.m. and 10 p.m. and show spatial as well as temporal consistency (at least over 10 years) in their acoustic features (Picciulin et al. 2012, Parmentier et al. 2017). If possible, based on signal-to-noise differences of simultaneously R-calling individuals, an approximation was made on the number of singers. In the presence of choruses, the abundance of calls is so high that individual call sequences cannot be discriminated any more (Fig. 1).

The large-scale sampling sites across the Northwestern Mediterranean were visually inspected for the presence of R-calls and choruses only, which are representative of courtship and spawning behaviour. On the other hand, long-term data was processed using a custom-made sound detection algorithm based on the rhythmic properties of calls and capable of discerning vocalizing individuals (Le Bot et al., 2015). This rhythmic analysis algorithm is based on time of arrivals of single pulses of a call within a frequency band of interest and uses a complex autocorrelation function with a sliding window to build a time-rhythm representation (Le Bot et al. 2015). The algorithm parameters were set to detect good-quality *S. umbra* calls that are generally temporally stereotyped (i.e., R-calls with mean pulse periods of 240ms; Parmentier et al. 2017) and with a minimum signal-to-noise-ratio of 6dB. Time series of the detected *S. umbra* calls were plotted over time. When available, temperature curves were provided and correlated with the acoustic activity.

## 3 RESULTS

### 3.1 Spatial distribution within MPAs

#### 3.1.1 Calanques National Park

The three recording sessions comprised a total of 72 recording stations covering over 40km of coastline and representing a surface of 20.4 km^2^ (considering a detection range of 300 m). 40 recording drifts were performed within NTZs (no-take zones) of the MPA and 32 in areas with a lower protection regime (RPZ) (Fig. 4). *S. umbra* R-& I-calls were recorded at 27.7% of the stations, both within (55% of call detections) and outside (45%) NTZs. R-calls (reproductive calls) equally occurred at 4 stations inside and 4 outside NTZ, indicating that reproductive activity also takes place in less protected areas. Most I-calls were detected inside NTZs (9 out of 12 stations). No choruses were detected and overall, calling activity was not abundant (only one, maximum two R-calls per stations and a few single I-calls), indicating a small number of vocalizing individuals.

**Figure 4.**
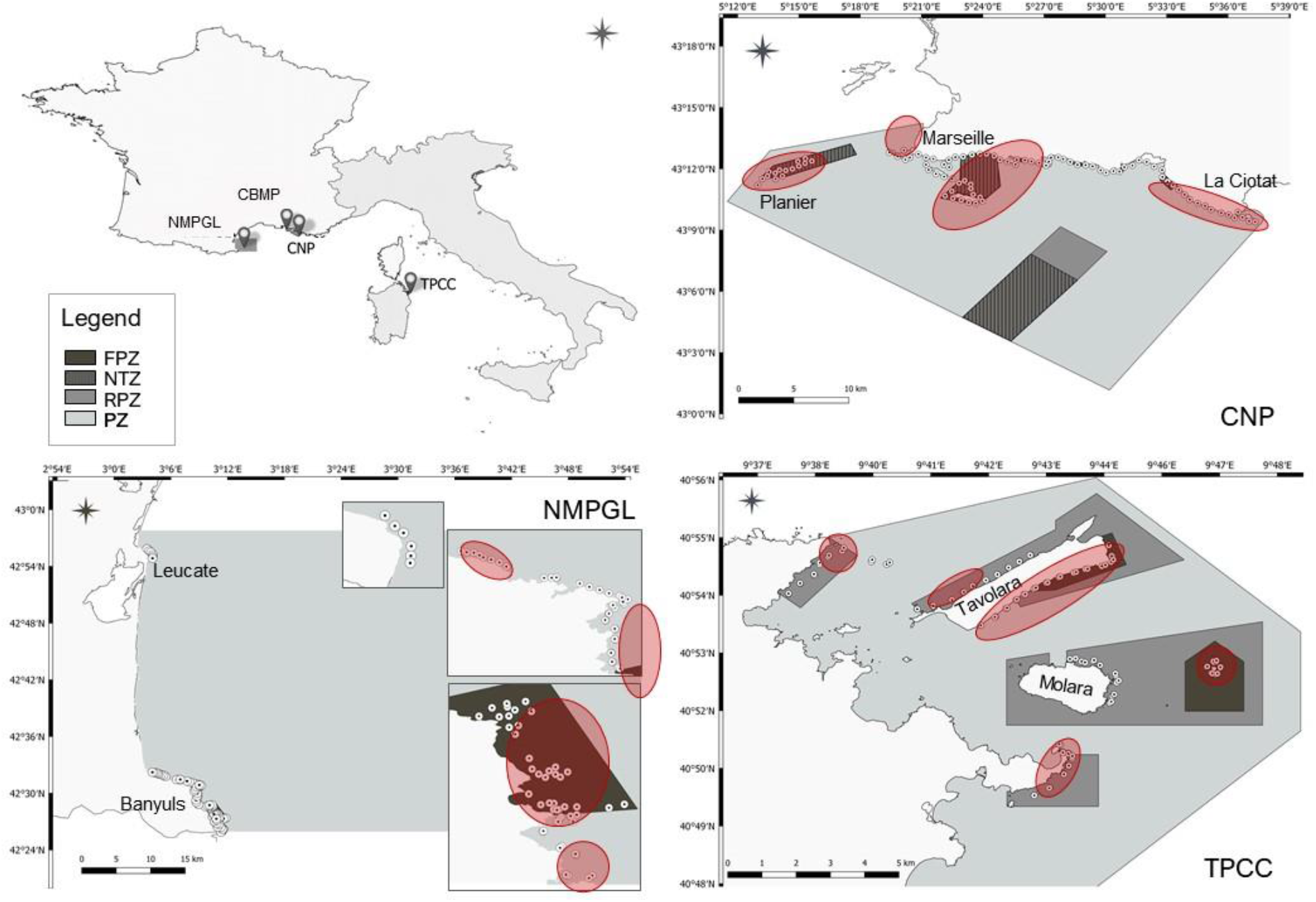
Maps of the drifting surveys and *S. umbra* detections within the MPAs. Top left panel: France and Italy with the locations of the four MPAs. Other panels: Maps of the three MPAs illustrating all the recording positions (white circles). Zones of *S. umbra* call detections are highlighted in red. Because of management and precautionary reasons with regard to illegal fishing, more detailed detection positions are not shown here. The different protection levels are indicated by different colours described in the legend: FPZ = Full Protection Zone (=Reserve = A zone), NTZ = No-Take-Zone (only in PNC), RPZ = Reinforced Protection Zone (= B zone), PZ = Protection Zone (= C zone). CNP: Calanques National Park, NMPGL: Natural Marine Park of the Gulf of Lion, TPCC: MPA of Tavolara-Punta Coda Cavallo.

#### 3.1.2 Marine Natural Park of the Gulf of Lion

96 sites were acoustically scanned during the four night-time surveys, allowing to cover the entire rocky coastline of the park (i.e., 30 km). *S. umbra* calls were present at 46 recording stations (i.e., 48% of the stations); 27 were within the marine reserve (MRCB) and 19 outside the reserve along the rocky cost of the marine park (Fig. 4). As the recording effort was greater within the reserve than in the rest of the park, there may be a bias towards the higher number of detections within the reserve. Nevertheless, the results show an effect of type and duration of the protection measures. In fact, the reserve was dense in detections, with *S. umbra* calls present in 77% compared to 33% of the stations in the rest of the park. Furthermore, in the reserve, acoustic reproductive behaviour was intense: 21 (of 27) stations with R-calls, three with choruses, and 50% of all stations with at least two singers. In the rest of the park outside the reserve, R-calls were recorded at 12 stations (66%), I-calls at 6 stations and simultaneous singing was detected at one station only. Overall vocal activity was therefore generally smaller compared to the reserve. From 4 sites recorded both in June and August in the NMPGL, two showed acoustic activity in both months, which may be indicative of site fidelity.

#### 3.1.2 MPA of Tavolara-Punta Coda Cavallo

54 stations were acoustically sampled during three recording sessions covering 20 km coastline: 16 in the full-protection zones (no-take & no-access) A, 24 in the RPZ zones B, and 14 in the less protected zone C (Fig. 4). At almost 70% of all stations, *S. umbra* calls were present. Zonation, i.e., the degree of protection and enforcement level, strongly influenced the observed calling behaviour: All stations in fully protected A zones showed sustained calling activity, including 11 stations with R-calls, one with I-calls and 2 with choruses. 12 out of the 16 stations showed elevated simultaneous singing of multiple individuals (2-4 or choruses). The B zone also showed a strong vocal activity (70% of the stations) with 10 stations in which R-calls were recorded, three in which I-calls occurred, three in which both R-and I-calls were present and one with a chorus. Simultaneous singing activity of at least 2 individuals occurred at 5 stations. The C-zone was the one with the fewest detections of *S. umbra* calls. Out of 14 stations, only 3 presented calling activity; one with I-calls, one with R-calls and one showing chorusing activity. Moreover, the Northern part of the MPA is more exposed to anthropogenic noise of ferries and other boats leaving the harbour of Olbia. Signal-to-noise ratio was overall small in N-Tavolara and Capo Ceraso because of increased background noise due to vessel traffic. Masking of communication signals may therefore be a cause for reduced signal production and/or detection.

### 3.2 Spatial sampling across the Northwestern Mediterranean basin

*S. umbra* was also present outside MPAs. On 23 out of 49 stations inspected, R-calls and choruses were detected (Fig. 5). This corresponds to an encounter probability of 47% (Fig 4) despite the fact that the recording stations were not specifically designed for *S. umbra* monitoring. The comparison between sites in seagrass meadows and coralligenous reefs indicates that *S. umbra* is acoustically present at 11 sites within seagrass meadows (50% of the sites) and at 8 sites on corallligenous reefs (30% of the sites). Although not quantified here, the number of simultaneously singing males was generally low (< 3 individuals), suggesting overall low fish densities. Potential spawning sites indicated by the presence of choruses only occurred at four sites; three in Corsica, and one in Sardinia. Two sites were within and two outside no-take zones.

**Figure 5.**
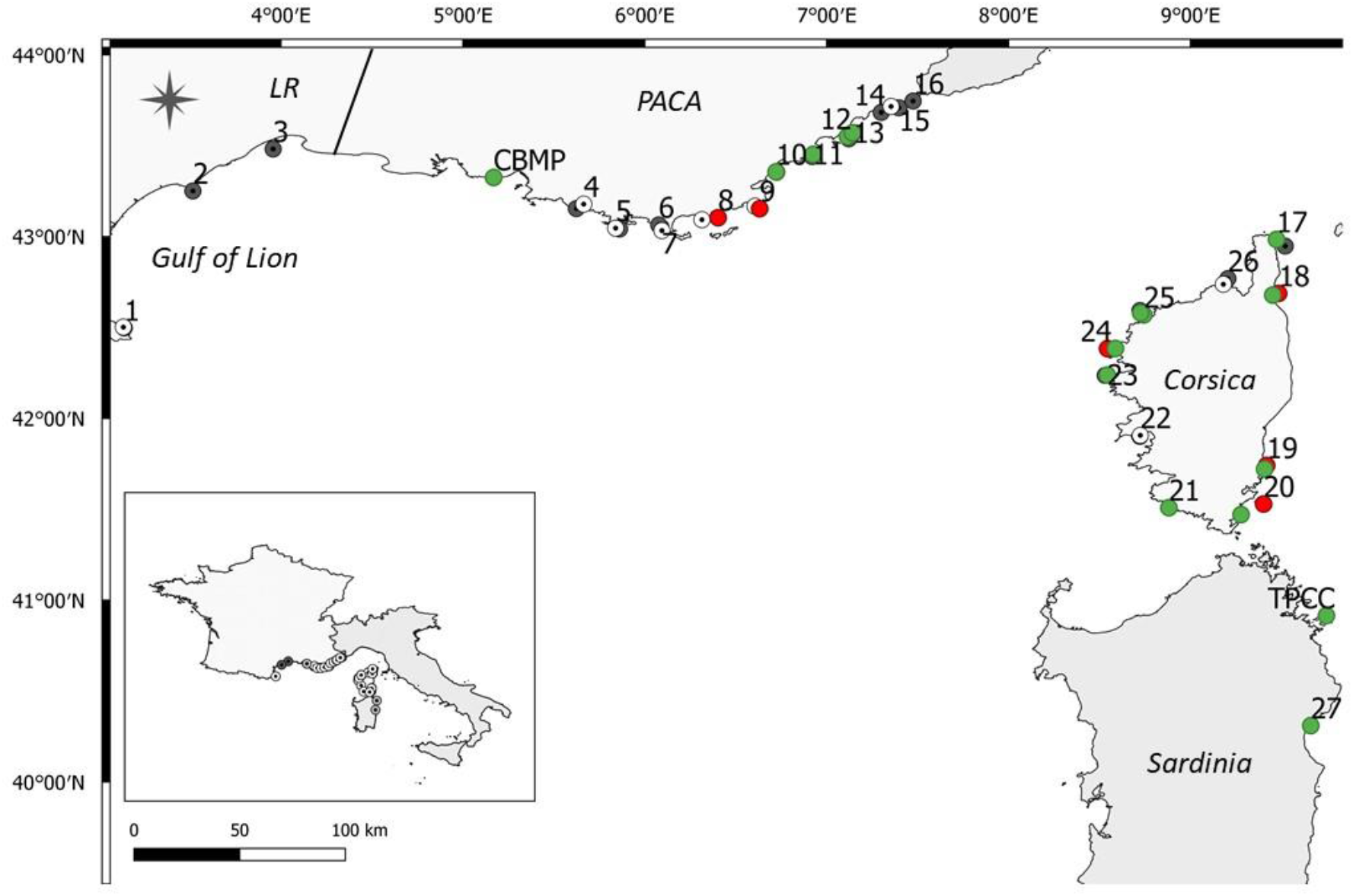
Map of the *S. umbra* detections within the acoustic monitoring network CALME. Green circles indicate *S. umbra* acoustic presence in the seagrass meadow and long-term recording sites, red circles indicate in the coralligenous reef sites. Dark grey circles indicate coralligenous reef recording sites without detections, and white circles are seagrass meadow recording sites without detections. CBMP: Côte Bleue Marine Park. *PACA*: Provence Alpes Côte d’Azur region, *LR*: Languedoc-Roussillon region.

### 3.3 Long-term seasonal monitoring within an MPA no-take zone

The annual recordings in the Reserve of Couronne in CBMP highlighted an acoustic activity of *S. umbra* spanning over a period of 5 months, with a strong consistency in sound production from one year to the other. In 2017, the first *S. umbra* calls were present in April but singing started on May 21 and stopped on October 11. In 2018 brown meagres started singing on May 22 and stopped on October 17 (Fig. 6). The singing period was almost identical and comprising part of spring and autumn. Singing activity was most intense during the summer months (June-September), and present 72% of the time (Fig. 6). Singing was also highly positively correlated with water temperature (ρ = 0.8) (Fig. 6).

**Figure 6.**
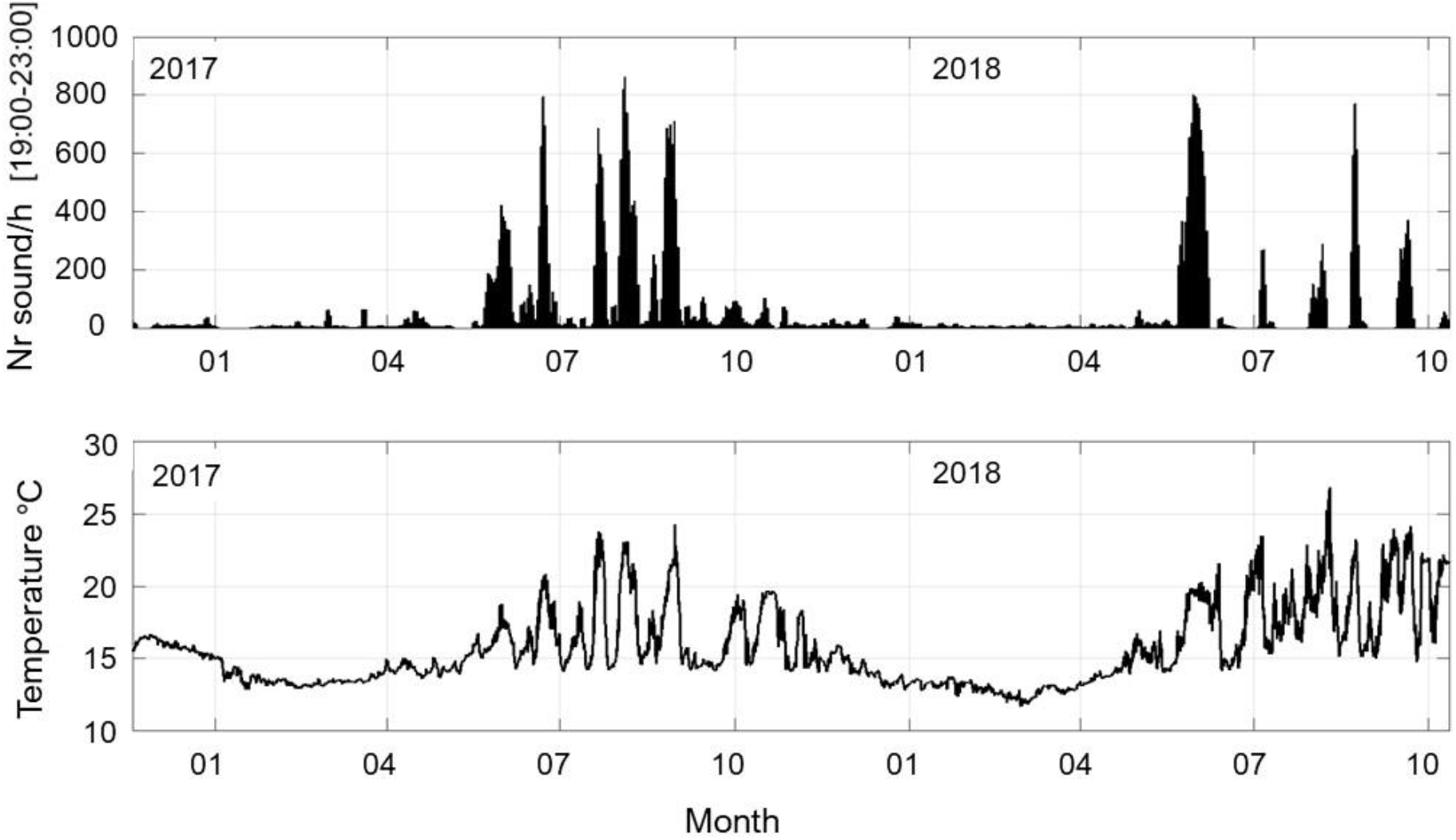
Biannual series of *S. umbra* calls in the Côte Bleue Marine Park, no-take Reserve of Couronne (top panel) and temperature profile at the recording site (bottom panel). Call production and temperature show strong correlations (ρ = 0.8).

## DISCUSSION

This study highlights the efficiency of PAM in studying and following *S. umbra* populations over different spatial and temporal scales for species management. The standardized method adopted to map the acoustic distribution of *S. umbra* using multiple drifting devices allowed to establish maps of the acoustic presence and reproductive activity of almost the totality of the MPAs’ rocky coastlines (up to 50km) in only 3-4 evenings. This confirms the efficiency of the method in rapidly assessing *S. umbra* distribution over vast areas (Picciulin et al., 2013). Distribution maps of this kind, obtained with limited human and logistic effort, are difficult to establish with traditionally used techniques. The results from these nocturnal surveys, showed that in the CNP and the NMPGL (excluding the reserve) *S. umbra* was present in approximately 30% of the recorded sites. In the Reserve of Cerbère-Banyuls (within the NMPGL) and TPCC, the probability of detecting *S. umbra* calls was between 70 and 80%, thus more than two times higher than in the NMPGL and the CNP. These differences may be linked to different management measures and particularly enforcement against anthropogenic pressures (i.e., fishing and diving), to the surveillance programs adopted by the different MPAs, to habitat characteristics, but also to the onset of the protection measures. In fact, the CNP and the NMPGL are young MPAs established approximately 7 years ago, while the MRCB and TPCC have been managed for over 20 years. Moreover, within the TPCC MPA, *S. umbra* calls were present in 100% of the sites within the fully protected zones A, 70% within reinforced B-zones and only 21% in the partially protected C-zones. Together these results indicate an overall effect of “age” of the MPA as well as a reserve-effect given by the level of protection and enforcement. This is consistent with the findings of long-term visual observations in Mediterranean MPAs indicating conspicuous increases in brown meagre abundance in fully protected or no-take zones over the last decade and an overall slow recovery of *S. umbra* populations (Francour, 1991; García-Rubies et al., 2013; Guidetti et al., 2014; Harmelin-Vivien et al., 2008; 2015; Harmelin and Ruitton, 2006). The role of enforced protection is further supported by the fact that almost all (8 out of 11) sites with chorusing activity, which are indicative of potential spawning aggregations, occurred within fully protected zones of older MPAs. Furthermore, in younger MPAs, calling activity was generally lower and the number of simultaneously singing individuals per site was smaller (< 3) compared to older more enforced MPA zones.

Overall, the R-call was the most abundant call type and it occurred within MPAs as well as in non-protected areas of the Northwestern Mediterranean. Outside MPAs, *S. umbra* reproductive calls were present in 47% of the sites, both in *Posidonia oceanica* seagrass meadows (50% of the seagrass sites) and in coralligenous reefs (30% of the coralligenous reefs). However, except for two chorusing sites in Corsica, the number of singers at one site was generally low, possibly indicating low animal densities. Corsica showed the highest number of sites with singing brown meagres (11 out of 23). On the mainland, they were mainly detected in the Eastern part of the region Provence-Alpes-Côte d’Azur (PACA, stations 8-6). In Western PACA only the CBMP reserve recorded brown meagre acoustic activity, while none was observed in the Languedoc-Roussillon region, which was however only represented by three stations (Fig. 5). These differences may be linked to the presence of suitable habitats for the species or fishing and may also be due to the fact that the recording sites were not specifically chosen to monitor the brown meagre. The coast of the gulf of Lion is dominated by sandy substrate with limited rocky and *P. oceanica* habitats. Overall, the *P. oceanica* sites with *S. umbra* detections were characterized by higher percentage of rocky substrate compared to the sites without detections (14% vs 6% rock). This may partly explain the distributions observed and why this species, typically associated to rocky habitats, was present in *P. oceanica* meadows. A more detailed analysis on habitat characteristics may provide insights to elucidate the observed distributions. The acoustic presence of *S. umbra* also outside MPAs, although generally in low densities, might be due to 1) the extended spatial coverage of the CALME network, 2) the presence of habitats suitable for the occurrence of the study species but also 3) the long-term management measures to increase water and habitat quality (EU Water Framework Directive) and 4) the ongoing moratorium in France that bans spearfishing and hook-and-line recreational fishing until December 2023 and thus partially protects *S. umbra* populations (Harmelin-Vivien et al., 2015; Rocklin et al., 2011).

Overall, these results show that, although strongly linked to habitat characteristics and protection level, *S. umbra* courtship activity occurs both within and outside MPAs (at least in France), and that PAM can be used to identify distribution and potential reproductive sites. Within the scope of spatial planning, PAM can contribute to define protection areas around functionally relevant zones (Erisman et al., 2012; Luczkovich et al., 2008b). This is particularly true for sites with chorusing activity that is associated with spawning aggregations, with chorus intensity being related to fish abundance and biomass in other Sciaenidds (Gannon and Gannon, 2010; Rowell et al., 2017). Large aggregations of the brown meagre have been reported in the Mediterranean Sea (Fiorentino et al., 2001; Ragonese et al., 2002). Surveys combining passive and active acoustics will help confirming the link between acoustic activity and fish abundance (Erisman and Rowell, 2017; Mann et al., 2008). Furthermore, in *S. umbra* it remains unknown if choruses are related to egg production and spawning as in the weakfish (*Cynoscion regalis*), spotted seatrout (*Cynoscion nebulosus*), red drum (*Sciaenops ocellatus*), silver perch (*Bairdiella chrysoura*), meagre (*Argyrosomus regius*) or the mulloway (*Argyrosomus japanicus*) (Lagardère and Mariani, 2006; Lowerre-Barbieri et al., 2008a; Luczkovich et al., 1999; Parsons et al., 2006). Ichthyoplankton and adult surveys at chorusing sites are required to assess the link between sound production and spawning (Luczkovich et al. 2008a). However, the results of the acoustic activity of *S. umbra* reported here clearly show that chorusing is rare (11 out of 271 sites) and prevails within fully protected or no-take zones in well-enforced MPAs. This is particularly relevant and further underlines the importance of marine reserves in protecting spawning sites and ultimately populations.

The vocal activity patterns reported here also allow to better describe reproductive movements of the brown meagre and the displacement of individuals for breeding aggregations that are still poorly documented (*c.f*., Harmelin-Vivien et al. 2015). PAM offers a means to identify the sites of vocal males and help assess the extent of their poorly known spatial movements and home ranges. It also shows its utility to fill some of the knowledge gaps in the nocturnal behaviour of this species (Harmelin-Vivien et al. 2015). *S. umbra* males in the Northwestern Mediterranean predominantly vocalize during the first half of the night, with a peak about one hour after sunset. This is therefore a time period at which, during the reproductive season, animals are engaged in reproductive behaviours. The diel pattern described here coincides with the one reported in Corsica (France), Sardinia (Italy) and in the Adriatic Sea (Italy) (Parmentier et al. 2017, Picciulin et al. 2013) and is commonly observed in many sciaenid species (Fish and Cummings, 1972; Mann and Grothues, 2009; Saucier and Baltz, 1993).

Two-year recordings in the Reserve of CBMP showed that the brown meagre is vocally active from May to October. Seasonal calling activity is common in Sciaenids (e.g., Connaughton and Taylor, 1994; Montie et al., 2015), but this is the first description of annual calling activity and natural variability in the brown meagre that may help better assess the spawning period of this vulnerable specie (Monczak et al., 2017). The spawning season of *S. umbra* is suggested to be from May to August, as estimated from histological examination and gonadosomatic index (Chauvet, 1991; Grau et al., 2009). Here we show that singing activity was sustained (i.e., 72% of occurrence) from late spring to autumn, indicating that the reproductive season of *S. umbra* is likely protracted, and occurring over a five-months period. Additional long-term recordings at different potential spawning sites will allow to confirm this extended reproductive season. The duration of the singing activity was equivalent in both years, starting on May 21 in 2017 and May 22 in 2018 and ending on October 11 in 2017 and October 17 in 2018. This relatively abrupt onset of calling activity suggests the presence of an environmental trigger. Temperature has been shown to initiate reproductive behaviour and chorusing in other sciaenid species (Mann and Grothues, 2009; Monczak et al., 2017; Rice et al., 2016). In this study, there was a strong positive correlation between water temperature and calling activity (ρ = 0.8), supporting the temperature-related hypothesis. Furthermore, although singing was sustained throughout the summer, it showed fluctuations in calling intensity (i.e., number of calls per hour). Peaks of vocal activity occurred in June, July, August and September and coincided with peaks in water temperature at the recording site. Similar temperature-related fluctuations in calling during the spawning season have been described in other Sciaenids (Connaughton and Taylor, 1994; Monczak et al., 2017; Montie et al., 2015) and have been related to seasonal cycles in the sonic muscle of males (Connaughton and Taylor, 1994). The smaller pulse-period observed during intense calling activity may be related to faster muscle twitch and contraction velocity at higher temperatures (Connaughton et al., 2002, Bolgan et al. 2020). An alternative explanation for the observed variations in calling intensity may be the influence of the lunar cycle (McCauley, 2012, Monczak et al., 2017). However, in this study intervals between peaks varied between 14 and 36 days (mean 25 days), thus indicating no clear lunar cycle pattern. In other sciaenid species, changes in calling behaviour have been related to spawning events (Lowerre-Barbieri et al., 2008; Mok and Gilmore, 1983; Montie et al., 2017). Although this remains to be elucidated for the brown meagre, the species is a multiple-spawning fish and within a population, females do not show synchronous ovarian maturity, which results in an extended spawning season (Grau et al. 2009).

## CONCLUSIONS

This study emphasizes the utility of PAM for the survey and conservation of exploited, vulnerable Mediterranean fish species. It represents a complementary, non-invasive, low-cost, standardized, replicable and efficient monitoring technique for environmental managers (Luczkovich et al., 2008a; Mann et al., 2008; Rountree et al., 2011). Here we focussed on the brown meagre, but the method can be extended to other vocal species (e.g., groupers). Harmelin-Vivien and co-authors (2015) underlined the need to combine extensive observations along the coast and regular long-term monitoring of favourable sites to document spatial distribution and the effect of increased protection of *S. umbra*. Here we show that PAM will allow to fill some of the knowledge gaps on species distribution, behaviour and reproduction needed for the conservation of the species using a combination of spatial and long-term acoustic recordings. Information of this kind is essential for the conservation of exploited or vulnerable fish species (Erisman et al., 2017).

The importance of recurrent monitoring surveys of threatened species inside and outside MPAs is critical to estimate population trends. Repeated standardized evening surveys over the season and across years will also allow, with limited human and logistic effort, to study site-fidelity and identify preferential sites of brown meagre populations over wide areas and help decision-makers to adopt protection acts and establish management zones (Luczkovich et al., 2008b). Here we showed that PAM increases our knowledge on the distribution and reproductive activity of this vulnerable species in rarely surveyed areas (i.e. outside MPAs) and over large geographical scales. PAM is also effective in emphasizing effects linked to management actions (e.g., no-take zones). Combining the obtained findings with existing visual observations, abundance estimations (e.g., using underwater visual census data or active acoustics), individual telemetry tracking data (Parsons et al., 2009; Taylor et al., 2006) as well as ichthyoplankton surveys on chorusing sites will provide critical information on the reproductive behaviour of the species, their distribution and population trends and allow to further consolidate the use of PAM for monitoring and managing this vulnerable species. Furthermore, simultaneous annual recordings on chorusing or known aggregation sites will help identify spawning events (e.g; Luczkovich et al., 2008b; Rice et al. 2015), trends in fish abundance via the calling activity (Erisman and Rowell, 2017; Rowell et al., 2017), and shifts in reproductive timelines associated with climate change (Monczak et al. 2017). Altogether, the findings reported here highlight the use of PAM as a means to support managers in their efforts to monitor, recover, and ultimately protect key ecological, commercial and emblematic species such as the Mediterranean brown meagre.

## CRediT authorship contribution statement

Lucia Di Iorio: Conceptualization, Methodology, Formal analysis, Writing – initial, review & editing, Investigation, Visualization, Supervision, Project administration. Patrick Bonhomme: Conceptualization, Investigation, Resources, Visualization, Funding acquisition. Noëmie Michez: Investigation, Resources, Writing - Review & Editing. Bruno Ferrari: Investigation, Resources. Alexandra Gigou: Conceptualization, Funding acquisition. Pieraugusto Panzalis: Investigation, Resources, Elena Desiderà: Investigation, Writing - Review & Editing. Augusto Navone: Resources, Pierre Boissery: Resources, Writing - Review & Editing, Funding acquisition. Julie Lossent: Methodology, Investigation. Benjamin Cadville: Investigation, Resources. Marie Bravo-Monin: Investigation, Resources. Eric Charbonnel: Resources, Writing - Review & Editing, Cédric Gervaise: Software, Methodology, Formal analysis, Conceptualization, Writing – review & editing, Investigation, Supervision, Project administration.

## ACKNOWLEDGEMENTS

The authors would like to thank the teams of the four MPAs as well as F. Cadène, V. Hartmann the team of the Marine Reserve of Cerbère-Banyuls for their technical support and their implication in the survey. Many thanks also to L. Bramanti, CNRS Observatoire Océanologique de Banyuls and the team of Andromède Océanologie for their support in the field. Thanks to J. Decelle for constructive comments on a previous version of the manuscript. This work has been possible thanks of the financial support of the Calanques National Park, the RMC Water Agency (convention 2016 1858), the French Agency for Biodiversity (AFB/2019 140), CHORUS Institute, and by the *Fonds Français pour l’Environnement Mondial* (FFEM) and the *Fondation du Prince Albert II de Monaco* via the MedPAN Calls for Small Projects 2019.

